# MetaRelSubNetVis: Referenceable network visualizations based on integrated patient data with group-wise comparison

**DOI:** 10.1101/2022.04.18.488628

**Authors:** Florian Auer, Simone Mayer, Frank Kramer

## Abstract

**Motivation:** Networks are a common data structure to describe relations among biological entities. Enriched with information to specify the entities or their connections, they provide a solid foundation for data-dependent visualization. When such annotations overlap, for example in a protein-protein interaction network that is enriched with patient-specific expressions, visualization is reliant on user interaction. Thereby, effective and reliable exchange of visualization parameters between collaborators is crucial to the communication within workflows.

**Results:** Here, we introduce MetaRelSubNetVis, a web-based tool that allows users to interactively apply group-wise visualizations to networks augmented with patient data. Our application can visually reflect patient-specific attributes for single patients or in a comparative context. Furthermore, we improved upon the exchange of network visualizations by providing unambiguous links that result in the same visual markup. Our work provides new prospects in interacting with and collaborating on network data, especially with respect to the exchange and integration of network visualizations.

## 1 Introduction

Networks are a well-established data structure in systems biology and often enhanced with annotations and integrated with additional data for their use in clinical applications (Heo et al., 2021). Visual exploration of these enriched networks is crucial for the interpretability of the contained information. Moreover, comparing different properties of a network, or even different networks based on the same properties remains an ongoing issue. Those networks may be composed of several patientspecific subnetworks based on preceding comparative analysis. An interactive investigation of these networks provides a more direct access to the information in contrast with static visualizations, the main purpose of which is mainly to communicate the results. Especially within a collaborative workflow individual investigations are required to gain necessary insights on the contained data. In turn this exchange of network data again requires specific infrastructure.

One well-established platform, where users can upload a network, share it with selected colleagues or make it publicly available, is the NDEx platform (Pratt et al., 2015). For stored networks, an interface is provided to integrate NDEx related services into third-party applications. One of those is Cytoscape (Shannon et al., 2003), a well-established network analysis and visualization software focusing on biological applications, and enables the exchange of networks with the platform. Cytoscape enables the visual exploration by defining mappings based on attributes of the integrated data but is impractical for quick changes between patient groups and properties. Furthermore, in a collaborative workflow the communication of the used visualization features is determining to the reproducibility and referenceability of the correct network visualization.

With MetaRelSubNetVis, we introduce a tool for the interactive group-wise visualization and comparison of integrated networks. In the following we elaborate on the example of patient-specific subnetworks, how our web-based application can facilitate the investigation of enriched networks in combination with the exchange of referenceable network visualizations.

## 2 Methods

MetaRelSubNetVis works with networks that are stored on the NDEx platform and can be referenced by its unique UUID and loaded in their proprietary Cytoscape Exchange (CX) format. CX is a JSON based data structure and was specifically designed for the data transmission. It originated from NDEx’s close connection to Cytoscape, where CX is used for the exchange of visualized networks.

The NDEx platform can be searched with MetaRelSubNetVis and the graph for the retrieved networks is rendered using the Cytoscape.js (Franz et al., 2016) library. The networks original layout and visualization are discarded during import and instead, a concentric layout is calculated and applied to provide a neutral visual markup. MetaRelSubNetVis is built upon Angular, an open-source framework for building single-page web-applications. The user interface was designed using Bootstrap, a well-established CSS-framework that provides a large set of front-end building blocks.

A CX network is composed of multiple aspects, each of which relates to a specific property of the network, for instance, nodes and edges, and accompanying attributes, meta information, layouts, and visual styles. Integrated data and annotations are conventionally found in the *nodeAttributes* and *edgeAttributes* aspects, while the description of the origin and composition of the data is contained within the *networkAttributes* aspect.

MetaRelSubNetVis requires detailed information about the enclosed integrated data to be able to use it for the creation of the data-dependent visualization and corresponding selectable options. This crucial information about patient data is stored within the *networkAttributes* aspect and involves their pseudonyms, group and subgroup affiliation, and for the exemplary network also patient survival details.

The web-application allows node coloring, sizing, and filtering based on the integrated data stored in the *nodeAttributes* for network wide and group wise properties. This can be for example the number of occurrences of relevant genes across all patients and the relevance scores for the genes in one specific patient, respectively. MetaRelSubNetVis relies on the definition of these visualization options within a for this purpose created non-standard aspect of the same name.

The formal requirements of the *metaRelSubNetVis* aspect are specified on the website, and additional scripts and documentation is provided as extension to the RCX library (Auer & Kramer, 2022) for the creation and handling within the statistical programming language R (R Development Core Team, 2008). The aspect allows the definition of continuous, discrete, and boolean mappings with their thresholds and corresponding color values. Furthermore, the included sample network hosted on the NDEx platform (UUID a420aaee-4be9-11ec-b3be-0ac135e8bacf) contains an implementation of the aspect and can provide guidance.

The sample network resulted from the analysis of a breast cancer data set for metastasis prediction and generation of patient-specific subnetworks by Chereda et al. Thereby, the complete protein-protein interaction network from the Human Protein Reference Database (HPRD) (Keshava Prasad et al., 2009) was used together with a large breast cancer dataset (Bayerlová et al., 2017) to predict for single patients the occurrence of a metastatic event and calculate a gene-wise score of its relevance for the prediction. The 140 most relevant genes of each patient were used to induce subnetworks, which were further combined to a single network and integrated with the gene expression values, levels, relevance scores, and subsequent Molecular Tumor Board report (MTB) analysis (Perera-Bel et al., 2018). MetaRelSubNetVis was then used for the investigation and visualization of the combined network within the original publication.

## 3 Results

### 3.1 Group-wise selection and network visualization

On the main page of MetaRelSubNetVis the user can select a network by searching the NDEx platform or continue with the provided sample network. The selected network is rendered with the default concentric layout and can be explored interactively by re-arranging its nodes. The position of the nodes will be kept even when the patient selection changes, or different layout setting are applied to facilitate the visual comparison of the subnetworks.

In the sidebar the user can adjust multiple settings (Fig. 1), while group-wise selection is one of the key aspects for this application. In the following the visualization options and its settings will be illustrated based on the provided sample network. In the patient dropdowns the user can select one sample per group for which the rendered network is updated with the corresponding subnetwork. Simultaneous selection of two patients leads to a comparative visualization in which the nodes are split and display the values according to the side of the group (Fig. 3).

**Fig. 1.**
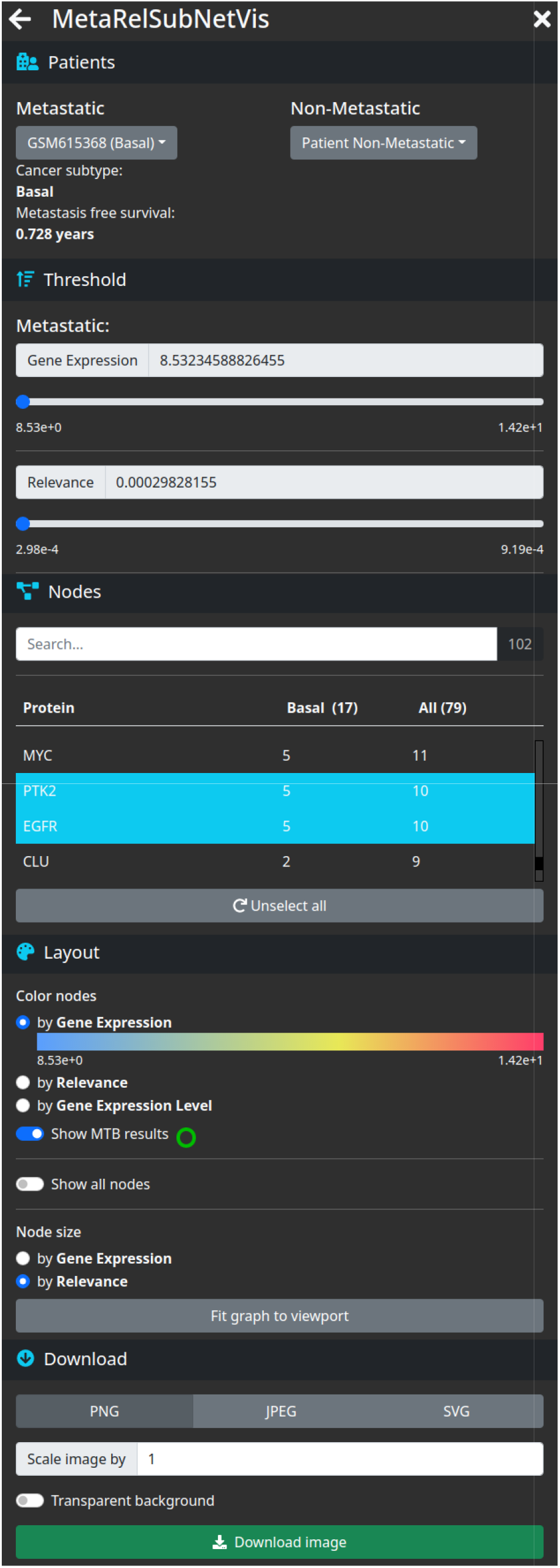
Selection and visualization options in MetaRelSubNetVis. For a selected patient the threshold can be adjusted, specific nodes selected, coloring and size of nodes c osen, and the visualization exported.

With a selected patient, the user can now adjust the thresholds of nodes to show. Adjusting this setting will hide nodes with lower values for those attributes. The nodes setting contains a list of all currently visible nodes and is searchable and interactive: Selecting nodes within the list will mark the respective node within the network. When one or two patients are selected, the list is augmented with information about the selected patient’s cancer subtype and occurrence of the genes.

The layout tab allows users to apply the in the *metaRelSubNetVis* aspect defined visualization options for the network. They can choose one of the predefined properties like gene expression level, gene expression or relevance score, to modify the coloring of the nodes. If only one patient is selected, they can adjust the size of each node with defined continuous mappings. Boolean properties as the MTB results allow to highlight corresponding nodes with a colored border.

MetaRelSubNetVis offers options to export the visualized network as image in three available data formats, namely PNG, JPEG and SVG. The image can be scaled with a factor up to 10 for non-SVG images and allows setting a transparent background for the PNG export.

### 3.2 Sharable visualization link

One significant aspect of MetaRelSubNetVis is the ability to quickly share network visualizations via a custom URL (Fig. 3). In the link generator tab users can highly customize the view, they want to share (Fig. 2). The table at the beginning provides a summary of the previously defined visualization options, such as the UUID of the network, selected patients, defined threshold, and marked and highlighted nodes.

**Fig. 2.**
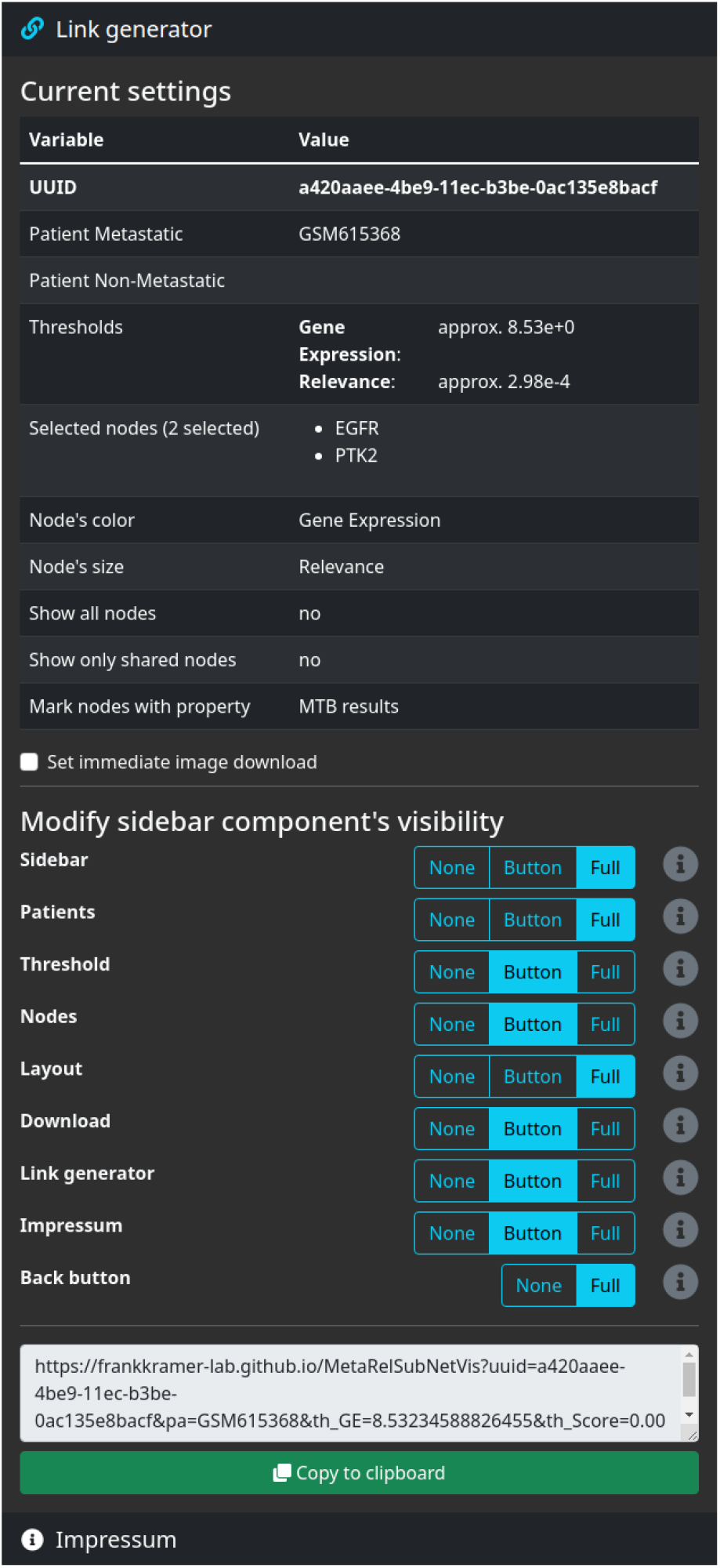
Options for creating a sharable link. Current settings for the visualization are listed and the behavior of the sidebar and its elements can be adjusted.

**Fig. 3.**
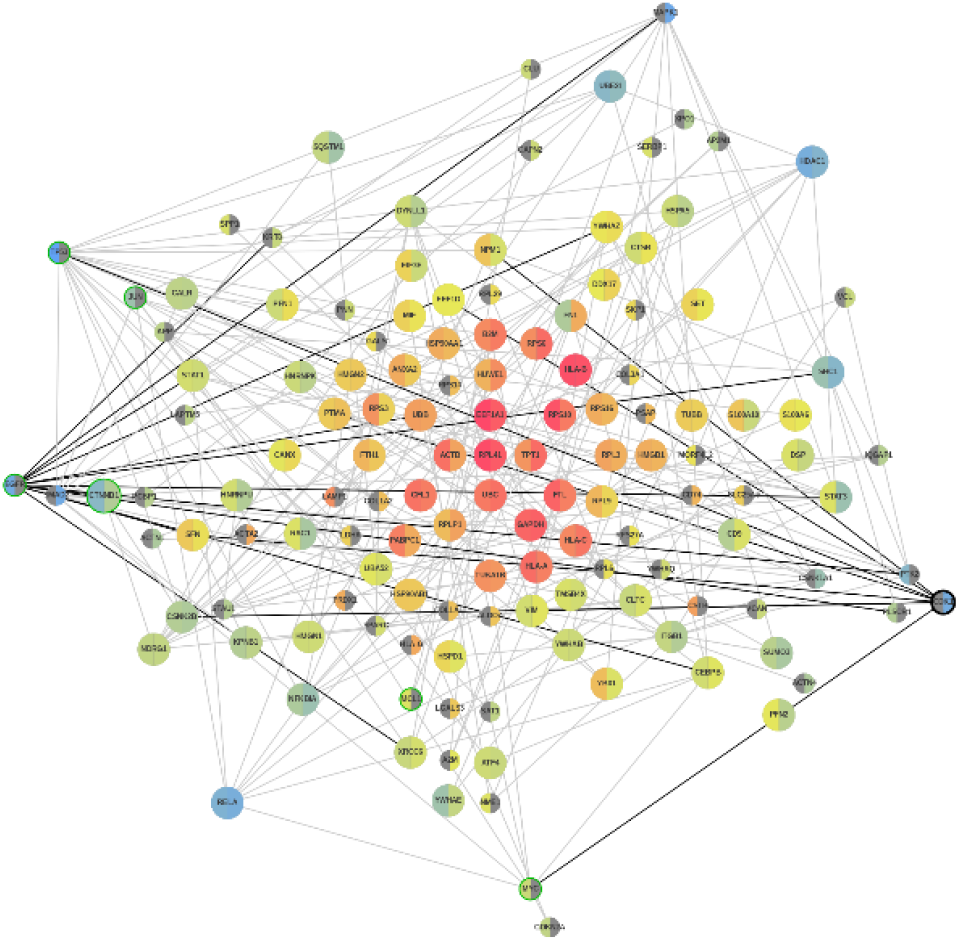
Visualization of gene expression in metastatic patients GSM615368 and GSM50093. The network visualization by MetaRelSubNetVis is available at https://frankkramer-lab.github.io/MetaRelSubNetVis?uuid=a420aaee-4be9-11ec-b3be-0ac135e8bacf&pa=GSM615368&pb=GSM50093&th_GE=8.53234588826455&th_Score =0.00029828155&sel=2319,3406&col=GE&size=Score&all=false&shared=false&bool= MTB&sb=0&cP=0&cT=1&cN=1&cL=0&cD=1&cG=1&cIm=1&bb=true

There may not be the need to inspect a network in the browser but rather continue working directly with an image of the network. In that case the user can decide to use the generated link to immediately trigger an image download in the specified format.

The rendering behavior of a network within MetaRelSubNetVis also includes customization of the sidebar. Each tab’s visibility, including the back button, and even the visibility of the whole sidebar can be defined in different stages. That opens exceptional possibilities, such as integrating a specific visualization of the network within an iFrame on a different web page. Hiding sidebar components that are not relevant or even the whole sidebar can be highly beneficial to direct the user’s focus on a particular aspect of the network.

## 4 Discussion

MetaRelSubNetVis was designed with focus on the visualization of a group-wise comparison between the integrated data. Yet, this approach is limited to two groups and for more than this it would also be difficult to be represented in an easily comprehensible manner. Additionally, the visualization is limited to the visualization of the same property within both groups. A potential extension could also be a comparative visualization of two properties for one selected patient. However, this would tremendously increase the dependency between visualization options and the required definitions for the visualization properties that it would be incompatible with the aim for simplicity of this application. On a related note, MetaRelSubNetVis only considers varying attributes relating to nodes. Patient-specific differences in edges are not yet respected and would be a challenge to render, especially for comparative visualizations. Splitting a node visually to show the expressions for the two respective patients is intuitively comprehensible, while a split or duplicated edge is hardly attributable to the corresponding node and may be perceived as confusing.

Providing information about the different groups and the data used for the integration of the network, as well as the definition of the visualization properties might be an obstacle to some users. However, the inclusion of meta-information should be a generally applied principle in network biology. Since the integration step already requires specialized knowledge, the effort required for the definition of the presumed visualization properties is negligible.

## 5 Conclusion

MetaRelSubNetVis is a web application that allows users to load a network enriched with group-specific information, inspect it and finally export or share the network. Retrieving the networks directly from NDEx not only promotes collaborative workflows through this platform but also circumvents the problems of finding individual hosting solutions for the used networks or incompatibilities due to different data formats.

Throughout the user’s visualization efforts, the positions of the single nodes remain consistent and thus improve comparability of the different enclosed properties of an integrated network. The group-wise comparison of network attributes allows a more comprehensible investigation of the results of preceding, already comparative analyses.

The communication and exchange of network visualizations is simplified with MetaRelSubNetVis by sharing a link to a specific layout configuration, facilitating collaboration furthermore. The options to hide specific parts of the sidebar or even the sidebar in total proves to be invaluable when embedding a network’s visualization within other web applications: Developers can highly customize the view without the implementation of an own proprietary network visualization.

MetaRelSubNetVis has already proven its potential by its application to the results of Chereda et al., where it was successfully used for the exploration, interpretation and visualization of the created specific subnetworks. patient-

## 6 Availability

A live version is hosted on GitHub Pages at https://frankkramerlab.github.io/MetaRelSubNetVis and the corresponding source code for deploying own instances is provided on https://github.com/frankkramerlab/MetaRelSubNetVis.

